# B7-H3 Modulates Cell Adhesion and Immune Evasion to Promote Tumor Progression and Natural Killer Cell Resistance in Hepatocellular Carcinoma

**DOI:** 10.64898/2026.03.28.714951

**Authors:** San Ha Han, Yoo Jin Cheon, Hyun Min Lee, Hyosun Seo, Ji Yoon Lee, Mi Jeong Kim, Suk Ran Yoon, Dongho Choi, Chun Jeih Ryu

## Abstract

B7-H3 (CD276) is an immune checkpoint molecule frequently overexpressed in hepatocellular carcinoma (HCC) and represents a promising therapeutic target. However, its roles in tumor cell adhesion, metastatic behavior and immune evasion—particularly in interactions with natural killer (NK) cells—remain incompletely understood. In the present study, B7-H3 was depleted using small interfering RNA and CRISPR/Cas9 in epithelial (Huh7 and HepG2) and mesenchymal (SNU449) HCC cell lines. Tumor cell survival, apoptosis, adhesion, migration and invasion were evaluated using functional assays. Expression of adhesion molecules and immune checkpoint proteins was assessed by flow cytometry and western blotting. Oncogenic signaling pathways were analyzed by examining phosphorylation of Akt, ERK, FAK and STAT3. NK cell-mediated cytotoxicity was assessed using primary human NK cells. B7-H3 depletion reduced clonogenic survival and increased apoptosis in mesenchymal HCC cells under anchorage-independent conditions. Loss of B7-H3 impaired cell adhesion, migration and invasion, accompanied by downregulation of PTGFRN, E-cadherin, integrin α3 and integrin αV, and reduced cell-to-cell aggregation under anchorage-independent conditions. B7-H3 depletion also decreased surface expression of PD-L1, PD-L2 and CD47. Notably, B7-H3-deficient cells exhibited enhanced susceptibility to primary NK cell-mediated cytotoxicity. Mechanistically, B7-H3 promoted tumorigenic signaling through Akt/S6, MVP/ERK and FAK/Src pathways in epithelial cells, and through FAK/Src and JAK2/STAT3 pathways in mesenchymal cells. Together, these findings reveal previously unrecognized roles for B7-H3 in coordinating adhesion and NK immune evasion in HCC, and support its therapeutic targeting for next-generation immunotherapies.

## Introduction

Hepatocellular carcinoma (HCC) is the most common form of primary liver cancer and a leading cause of cancer-related mortality worldwide [1]. Although long-term survival is possible in some HCC patients by surgery, trans-arterial chemoembolization and transplantation [2], HCC has high recurrence rate (> 70%) due to metastases [3, 4]. The complex and aggressive nature of HCC, often accompanied by a poor prognosis, highlights the urgent need for new therapeutic strategies. Tyrosine kinase inhibitors (TKIs, Sorafenib, Lenvatinib) are used in patients with HCC that have metastasized or recurred, but the survival benefit of TKIs is only extended by a maximum of 13.6 months [5]. A key feature of many cancers, including HCC, is their ability to evade the host immune system through mechanisms involving immune checkpoint molecules [6]. Recently, several systemic therapies, including immune checkpoint inhibitors (ICIs) such as Atezolizumab and Bevacizumab, have received FDA approval for the treatment of advanced HCC [1, 7]. The combination therapy of Nivolumab and Ipilimumab has also been approved by the FDA [8]. The outcomes of these therapies have shown a significantly improved survival benefit [1, 8]. However, objective response rates are around 30% and a significant portion of HCC patients do not respond to existing immunotherapies [1, 8], suggesting that other checkpoint molecules or evasion pathways are involved.

B7-H3 (CD276) is an immune checkpoint protein that is highly expressed in various solid tumors but shows minimal expression in normal tissues, making it an attractive pan-cancer therapeutic target [9]. Many studies have shown various non-immunological actions of B7-H3 in cancer development. B7-H3 is linked to cancer cell proliferation, anti-apoptosis, pro-angiogenesis, metastasis, metabolic reprogramming and therapeutic resistance [10]. B7-H3 is also involved in the immunosuppression of tumor cells by reducing interferon secreted by T cells and inhibiting the cytotoxic activity of natural killer (NK) cells [11, 12]. Previous studies, including our own, have shown that B7-H3 is involved in the proliferation, migration, and invasion of cancer cells [12, 13]. We have previously demonstrated that a monoclonal antibody, NPB40, which targets B7-H3, significantly inhibits tumor growth in an HCC xenograft mouse model [13]. These findings strongly suggest a critical role for B7-H3 in HCC progression in vivo, but the specific cellular and molecular mechanisms underlying this role remain to be fully elucidated. B7-H3 is overexpressed (79-94% positivity) in HCC tissues and is closely associated with HCC aggressiveness, recurrence and poor survival [14]. This study aims to investigate the multifaceted functions of B7-H3 in HCC cells. We explored its impact on key cellular processes, including survival, apoptosis, adhesion, migration, and invasion. Specially, we focused on its relationship with cell adhesion molecules and immune checkpoint molecules, and analyzed its role in regulating NK cell-mediated cytotoxicity. Furthermore, we sought to determine the specific signaling pathways through which B7-H3 exerts its effects in different HCC cell types (epithelial and mesenchymal). This study identifies a novel role for B7-H3, suggesting that B7-H3 comprehensively regulates cell adhesion and immune evasion of HCC cells and provides strong evidence for the development of B7-H3 as an excellent therapeutic target for the treatment of HCC.

## Materials and methods

### Cell culture

Human cancer cell lines Huh7 and SNU449 were purchased from the Korean Cell Line Bank (KCLB, Seoul, Korea) and cultured in RPMI-1640 (Biowest, Kansas, MO, USA) medium supplemented with 10% fetal bovine serum (FBS, Corning, Seoul, Korea) and antibiotic-antimycotic solution (AA, Biowest). HepG2 cells were purchased from American Type Culture Collection (ATCC, Manassas, Virginia, USA) and maintained in MEM (Corning) medium supplemented with 10% FBS (Biowest) and AA solution (Biowest). HEK293FT cell line (Thermo Fisher Scientific, Waltham, MA, USA) was cultured in DMEM (Welgene, Gyeongsan, Korea) medium supplemented with 10% FBS and AA solution. The NK92 cell line was purchased from ATCC and cultured in α-MEM (Biowest) medium supplemented with 12.5% FBS (Biowest), 12.5% horse serum (Biowest), 200μM inositol (Sigma-Aldrich, St.Lois, MO, USA), 20μM Folic acid (Sigma-Aldrich), 100μM 2-mercaptoethanol (Sigma-Aldrich), AA solution (Biowest), and 400 unit/ml IL2 (Peprotech, Rocky Hill, NJ, USA). Cancer cells were cultured according to the instructions provided by the suppliers.

### Western blotting

For general Western blotting, cells were lysed with RIPA lysis buffer (25 mM Tris-HCl pH 7.6, 150 mM NaCl, 1% NP-40, 1% sodium deoxycholate, 0.1% SDS) and denatured by boiling with sample buffer. Protein samples were separated by 10% SDS-PAGE gel for 100 min, and the separated proteins were transferred to a nitrocellulose membrane for 1 hour 30 min at 80V. A commercial protein ladder (Themo Fisher Scientific #26616) was used to analyze protein size. The membranes were blocked with blocking buffer (5% skim milk with Tris-buffered saline with 0.1% Tween 20 (TBST)) for 2 hours at room temperature (RT) and washed with 0.1% TBST 3 times. Blocked membranes were incubated with primary antibodies (0.1∼2 μg/ml) diluted with blocking buffer overnight at 4°C. Primary and secondary antibodies used in his study were described in **S1 Table**. For detection of the phospho-form of proteins, membranes were incubated with primary antibodies diluted with blocking buffer. The membranes were then washed with 0.1% TBST 3 times and incubated with goat anti-mouse IgG-HRP or goat anti-rabbit IgG HRP for 1 hour at RT. The membranes were analyzed with Western bright ECL HRP substrate (Advansta, Menlo Park, CA, USA).

### Mass Spectrometry

Immunoprecipitants were separated on 8% polyacrylamide gel and stained with PageBlue™ Protein Staining Solution (Thermo Fisher Scientific). To identify proteins, LC-MS/MS analysis was performed at ProteomeTech (Seoul, Korea). The search program ProFound was used for protein identification [15].

### siRNA and transfection

Small interfering oligonucleotides targeting B7-H3 (siB7-H3; Bioneer, Daejeon, Korea) were always used with control siRNA (siCon, Genolution, Seoul, Korea). Cells were transfected with 50 nM siRNA by using RNAiMAX (Thermo Fisher Scientific) following the manufacturer’s protocol. The cells were transfected the second time after 24 hours following the first transfection and then incubated for additional 48 hours before harvesting for analysis.

### Generation of B7-H3 knockout (KO) cell lines

The guide sequence oligonucleotides (CACCGTTCGTGAGCATCCGGGATTT and AAACAAATCCCGGATG CTCACGAAC), including BsmBI restriction site overhangs and targeting exon 5 of B7-H3 [16], were annealed and cloned into the lentiCRISPRv2 vector (Addgene #98290, Watertown, MA, USA). To generate lentiviral particles, HEK293FT cells were seeded at a density of 3×10⁶ cells per 100-mm culture dish one day prior to transfection. Cells were transfected with 3 μg of psPAX2 (Addgene #12260), 1 μg of pMD2.G (Addgene #12259), and 5 μg of the transfer vector in 600 μl of serum-free medium containing 27 μl of PolyJet™ (SignaGen, Frederick, MD, USA). Plates were incubated at 37°C in a humidified CO₂ incubator, and the culture medium was replaced 24 hours after transfection. Lentiviral supernatants were collected at 48 and 72 hours post-transfection and filtered through a 0.45-μm Millex sterile syringe filter (Millipore, Burlington, MA, USA). Serial dilutions of the lentiviral suspension supplemented with Polybrene (8 μg/ml; Sigma-Aldrich) were added to 2×10⁵ SNU449 cells cultured in 1 ml of fresh medium. After 24 hours, the medium was replaced with fresh culture medium containing puromycin (InvivoGen, San Diego, CA, USA). Single-cell clones were obtained by serial dilution in 96-well plates, and B7-H3 expression was assessed by Western blotting and flow cytometry.

### Quantitative polymerase chain reaction (qPCR)

Total RNAs were isolated from control SNU449 and B7-H3 KO SNU449 using the RNAiso Plus (Takara, Seoul, Korea) according to the manufacturer’s instructions. RNAs were converted to cDNAs using PrimeScript^TM^ RT reagent Kit (Takara). qPCR was performed using Applied^TM^ Biosystem PowerTrack^TM^ SYBR Green PCR Master Mix (Thermo Fisher Scientific). The target gene-specific primers for human transcripts encoding B7-H3 [17], PTGFRN, E-cadherin, ITGAV, ITGA3, GAPDH are listed in **S2 Table**. Relative expression levels between different samples were calculated using ΔΔCt method [18].

### Apoptosis assays

To detect apoptosis, control and B7-H3 knockdown HCC cells were cultured for 3 days and stained with propidium iodide (PI) and fluorescein isothiocyanate (FITC)-conjugated annexin V (BD Biosciences, Seoul, Korea) according to the manufacturer’s protocol. For apoptosis induction, cells (5 × 10^5^ cells) were seeded in ultra-low attachment 6-well plates (Corning) with serum-free media. Plates were incubated for 2 days at 37°C in a humidified CO_2_ incubator. After incubation, cells were stained with PI and FITC-conjugated annexin V (BD Biosciences). The cells were also lysed with RIPA lysis buffer and used for the Western blotting. The stained cells were analyzed by FACS Canto (BD Biosciences) with the FACS Diva software.

### Clonogenic survival, cell migration, and invasion assays

Cells (0.1-1 × 10^4^ cells) were plated on 6-well plates with complete medium. After 6-8 days of incubation, colonies were fixed with 4% paraformaldehyde (PFA) solution (Sigma-Aldrich) and stained with 0.5% crystal violet solution (Sigma-Aldrich) overnight at RT. The stained colony area and the intensity were measured using the Image-J program. Migration and invasion assays were performed in Trans-well chambers (Corning) as described previously [19].

### Cell adhesion, spreading, and aggregation assays

For cell-adhesion assays, twelve-well plates were coated with gelatin for at least 1 hour at 37℃. HCC cells were seeded at 1 × 10^6^ cells per well and incubated for 3 hours at 37℃. The plates were washed with phosphate buffered saline (PBS, pH 7.4) to remove unattached cells. Adherent cells were stained with 0.5% crystal violet solution for 30 minutes at RT. The stained cells were counted in photographic image after washing with distilled water, or dissolved in 0.1% SDS before measuring the optical density (OD) at 570 nm. For cell spreading assays, cells were seeded with 5 × 10^4^ cells in 1 ml culture media per well in the Matrigel (Corning)-, gelatin (Sigma-Aldrich)- or collagen IV (Thermo Fisher Scientific)-coated 12-well plates and incubated for 1 hour at 37°C in a humidified CO_2_ incubator. After incubation, the attached cells were fixed with 4% PFA and incubated for 15 min at RT for fixation. Next, fixed cells were carefully washed with PBS (pH 7.4) 2 times, and observed the cell spreading morphologies using the Image-J cell spreading plug-in. For cell aggregation assays, cells were resuspended through a 0.44 μm cell strainer (SPL, Pocheon, Korea), then single cell suspensions were seeded into 6-well ultra-low attachment plates (Corning) for 5 × 10^4^ cells per well. The plates were incubated for 1 hour at 37°C in a humidified CO_2_ incubator with shaking. Before and after incubation, single and contact cell numbers were counted, and the cell aggregation percentage was measured.

### Differentiation of primary NK cells from umbilical cord blood (UCB)

Human UCB was obtained from healthy pregnant woman with consent and the study was conducted with the procedures approved by the Institutional Review Board (IRB) at Korea Research Institute of Bioscience and Biotechnology (IRB No. P-01–201610-31–002). Primary NK cells were differentiated from human UCB cells as described previously (Lee, Park et al. 2022). Briefly, human UCB cells were first depleted T cells and red blood cells using Rosette Sep (StemCell Technologies, Vancouver, Canada) and the mononuclear cells (MNC) were isolated by Ficoll-Hypaque density centrifugation according to the manufacturer’s instructions. For in vitro NK cell differentiation, CD3+ T cell-depleted MNCs were cultured in α minimum essential medium (Welgene) supplemented with 10% FBS (Themo Fisher Scientific), 1% penicillin/streptomycin (Gibco), 10^-6^ M of hydrocortisone (Stem Cell Technologies), IL-15 (10 ng/ml) and IL-21 (10 ng/ml). The cells were cultured for 10 days and half of the medium was replaced every 2 days supplemented with fresh cytokines. All cytokines for NK cell differentiation were purchased from PeproTech.

### NK cell-mediated cytotoxicity assays

For NK cell-mediated cytotoxicity assays, target cells were seeded in 12-well plates at 2 × 10^4^ cells per well. The plates were incubated overnight at 37°C in a humidified CO_2_ incubator. After incubation, the culture media was changed, and NK92 cell suspension was added to each well at a 1:1 E:T ratio and incubated for 2 days at 37°C in a humidified CO_2_ incubator. The plates were carefully washed with PBS (pH 7.4) 2 times, and the cells were fixed with 4% PFA and incubated for 15 min at RT for fixation. After fixation, the fixed cells were washed with PBS (pH 7.4) 2 times and stained with 0.5% crystal violet solution for overnight at RT. The stained cells were washed with distilled water to completely remove the remaining solution and then dried at RT. The stained cell density was measured by dissolving the stained cells with 10% acetic acid solution and measuring the OD values at 590 nm. For calcein-acetoxymethyl (calcein-AM) cytotoxicity assay, 1 × 10^6^ target cells were washed and resuspend with serum-free media containing 3 μg/ml calcein-AM reagent (Biolegend, San Diego, CA, USA) and incubated for 30 min at 37°C in a humidified CO_2_ incubator. Target cells were then washed with PBS (pH7.4) twice and were resuspended in NK92 cell media at a concentration of 5 × 10^5^ cells/ml. NK92 or primary NK cells were added to target cells at various ratios (0.5:1∼1:10, target cells: effector cells with all samples in triplicate) in the presence of IL2 (400U/ml). Each 100 μl of target and effector cell suspensions was then centrifuged for 3 min at 400 × g in 96 well round bottomed plates and incubated for 5 hours at 37°C in a humidified CO_2_ incubator. For the calcein-AM release assay, 150 μl of the supernatant was collected and transferred to a flat bottom black plate. The fluorescence was read using a BioTek Synergy 2 plate reader (Ex: 485 nm / Em: 535 nm). The percent specific lysis was calculated using the formula [(test release-spontaneous release)/(maximum release-spontaneous release)] × 100.

### Statistical analyses

Statistical evaluation for data analysis was performed using GraphPad Prism software and was determined by unpaired Student’s t-test or two-way ANOVA. All experiments were performed at least three times. All data are presented as mean values ± standard deviation (SD). A p-value <0.05 is indicated by *, p <0.01 by **, and p <0.001 by ***.

## Results

### B7-H3 promotes clonogenic survival and apoptosis resistance in HCC cells

Previously, we generated NPB40, a monoclonal antibody targeting B7-H3 expressed in human embryonic stem cells, and demonstrated that B7-H3 acts as a potential cancer stem cell marker in HCC cells, and NPB40 injection significantly inhibits tumor growth in an HCC xenograft mouse model [13]. These results suggest that B7-H3 plays a critical role in the progression of HCC in vivo. To further investigate the role of B7-H3 in HCC cells, *B7-H3* was transiently depleted by siRNA technology in HCC cells. We used three different HCC cell lines, in which Huh7 and HepG2 belong to epithelial HCC cells and SNU449 belongs to mesenchymal HCC cells [20]. HCC cells were transfected with either negative control siRNA (siCon) or *B7-H3* siRNA (siB7-H3). *B7-H3* depletion decreased the numbers and sizes of Huh7 and HepG2 colonies by approximately 3.6- and 1.4-fold, respectively [13]. To examine whether B7-H3 depletion also induces apoptosis in epithelial HCC cell lines, *B7-H3*-depleted Huh7 cells were stained with annexin V and PI. Single PI-positive and both annexin V and PI-positive cells were increased by approximately 10% in *B7-H3*-depleted Huh7 cells (**Fig. 1A, 1B**). *B7-H3* depletion also induced decreased expression of BCL-xL in Huh7 cells, whereas it increased the expression of Bax (**Fig. 1C**). Single PI-positive HepG2 cells were also increased by approximately 11% in *B7-H3*-depleted HepG2 cells (**Fig. 1D, 1E**). The increased expression of BAX was also observed in *B7-H3*-depleted HepG2 cells, although the expression of BCL-xL was not significantly altered (**Fig. 1F**). The results suggest that B7-H3 promotes cell survival in epithelial HCC cells by inducing survival factor and inhibiting apoptosis regulator.

**Fig. 1.**
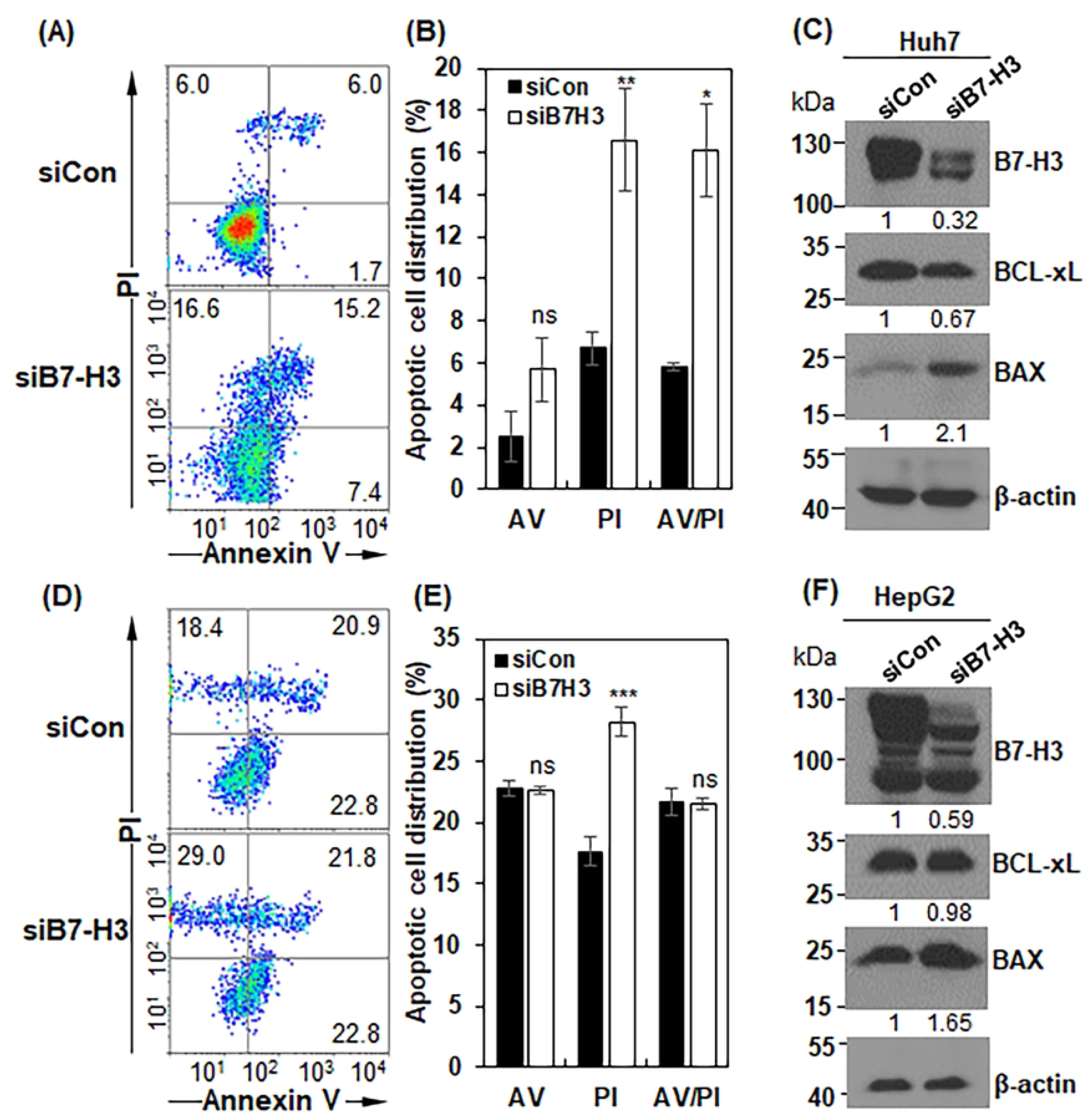
B7-H3 knockdown enhances apoptosis in Huh7 and HepG2 cells. (A) Apoptosis was analyzed by Annexin V and propidium iodide (PI) staining in B7-H3–knockdown Huh7 cells. (B) Quantification of apoptotic Huh7 cells shown in (A). (C) Expression of apoptosis-related proteins in control and B7-H3–knockdown Huh7 cells was analyzed by Western blotting and quantified using ImageJ software. (D) Apoptosis analysis by Annexin V/PI staining in B7-H3–knockdown HepG2 cells. (E) Quantification of apoptotic HepG2 cells shown in (D).

To investigate the role of B7-H3 in mesenchymal HCC cells, *B7-H3* was also transiently depleted by siRNA technology in SNU449 cells. B7-H3 protein was decreased approximately 93% in si*B7-H3*-transfected SNU449 cells (**S1A Fig**). When cell surface B7-H3 was measured with NPB40 and polyclonal anti-B7-H3 antibodies (α-B7-H3), the expression levels decreased to approximately 91% in si*B7-H3*-transfected SNU449 cells (**S1B-1C Fig**), demonstrating that si*B7-H3* sufficiently depletes B7-H3 expression in SNU449 cells. B7-H3 depletion also decreased the numbers and sizes of SNU449 colonies by approximately 3.2-fold (**Fig. 2A**). To further investigate the role of B7-H3 in SNU449 cells, *B7-H3* was also permanently depleted by CRISPR/Cas9 technology with *B7-H3* targeting sgRNA. Western blotting showed that B7-H3 proteins were almost depleted in 5 subclones except for A11 subclone (**S1D Fig**). Flow cytometry analysis also showed that cell surface expression of B7-H3 was drastically suppressed in the 5 KO subclones (**S2E, S2F Fig**). A representative subclone, B3, showing drastic reduction in B7-H3 expression was selected for further experiments (**S1G-S1J Fig**). Then, clonogenic survival assays were performed with the representative *B7-H3* KO subclone and *B7-H3* KO decreased the colony areas of SNU449 colonies by approximately 1.8-fold (**Fig. 2B**). The results indicate that B7-H3 enhances clonogenic survival and proliferation of SNU449 cells. To examine the role of B7-H3 in apoptosis in SNU449 cells, *B7-H3* KO SNU449 cells were stained with annexin V and PI. Contrary to epithelial HCC cells, however, annexin V-and annexin V/PI-positive cells were not significantly changed in adherent *B7-H3* KO SNU449 cells (**Fig. 2C, 2D**). To induce apoptosis, therefore, SNU449 cells were seeded in ultra-low attachment 6-well plates with serum-free medium. When apoptosis was induced by culturing the cells in an anchorage-independent manner, annexin V-positive cells were increased by approximately 24% in *B7-H3* KO SNU449 cells, as compared to those of control cells (**Fig. 2C, 2D**). These findings suggest that B7-H3 promotes clonogenic survival and inhibits apoptosis in HCC cells. Specifically, B7-H3 can promote apoptosis resistance in mesenchymal HCC cells even under anchorage-independent conditions.

**Fig. 2.**
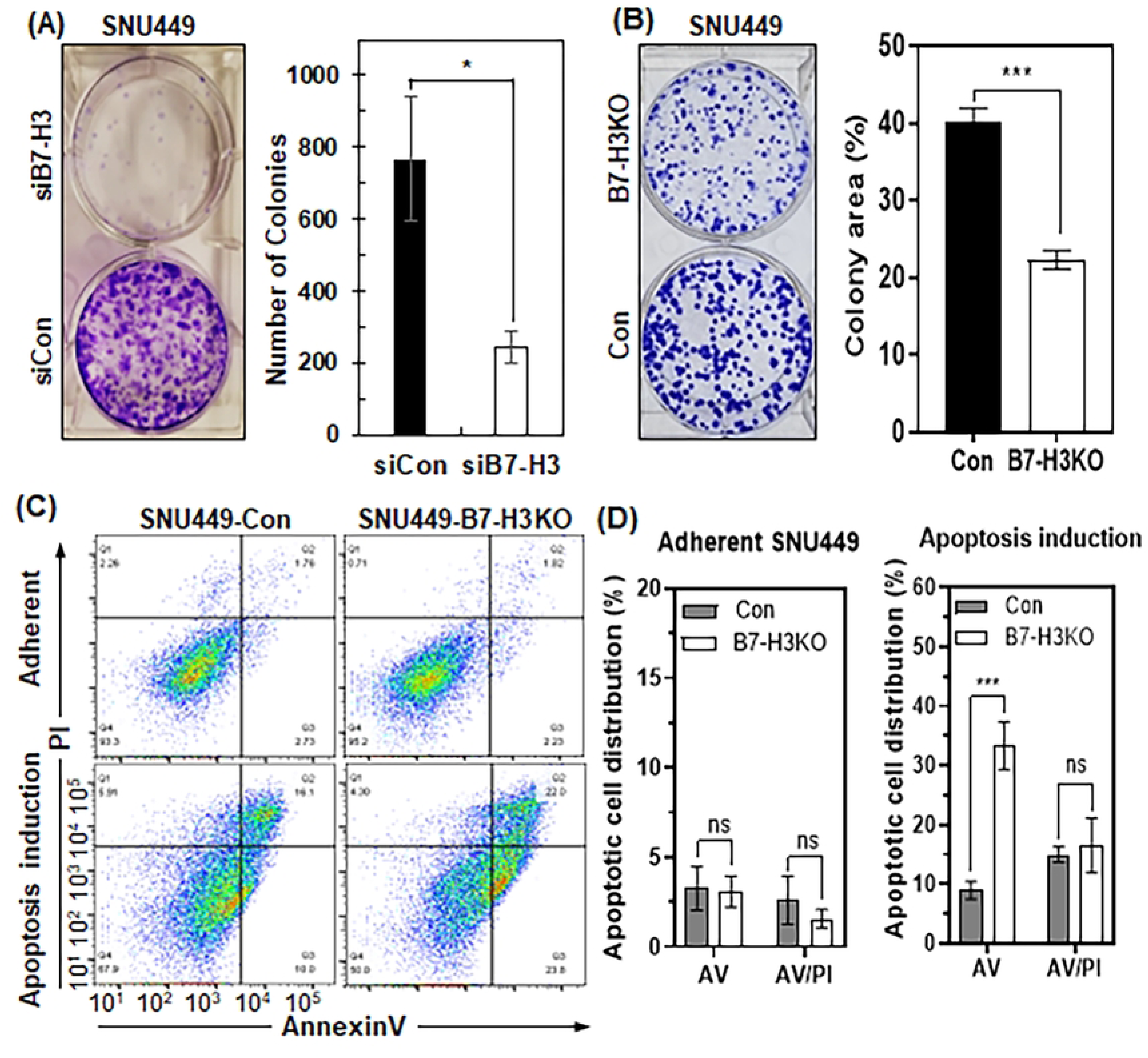
B7-H3 promotes clonogenic survival and apoptosis resistance in HCC cells. (A) Clonogenic survival assay of SNU449 cells transfected with control siRNA (siCon) or B7-H3 siRNA (siB7-H3). Colony numbers were quantified using ImageJ software. (B) Clonogenic survival assay of control and B7-H3 knockout (KO) SNU449 cells. Colony areas were quantified using ImageJ software. (C) Apoptosis analysis by Annexin V/PI staining in control and B7-H3 KO SNU449 cells under adherent conditions and after apoptosis induction under anchorage-independent conditions. (D) Quantification of apoptotic cells shown in (C). ns, not significant; *, p < 0.05; ***, p < 0.005.

### B7-H3 promotes cell adhesion, migration and invasion

Cell adhesion plays an important role in cancer cell growth, survival and metastasis [21]. To examine whether HCC cell adhesion to extracellular matrix (ECM) is affected by *B7-H3* depletion, *B7-H3*-depleted Huh7 cells were incubated with gelatin-coated plates, and the number of adherent Huh7 cells was quantified by the OD of crystal violet-stained cells (**Fig. 3A, 3B**). Adherent Huh7 cells were decreased by 33% in *B7-H3*-depleted Huh7 cells. The same cell adhesion assays were also done with HepG2 cells, and adherent HepG2 cells were decreased by 25% in *B7-H3*-depleted cells (**Fig. 3C, 3D**), suggesting that B7-H3 expression promotes HCC cell adhesion to ECM. To study how B7-H3 regulates cell adhesion, we examined the cell surface expression of cadherin and integrin molecules by flow cytometry in *B7-H3*-depleted Huh7 cells (**Fig. 3E, 3F**). Cell surface expression of PTGFRN (CD315), E-cadherin, integrin α3 and integrin αV was decreased by approximately 84%, 68%, 25% and 17%, respectively, in *B7-H3*-depleted Huh7 cells. Similar results were also observed with HepG2 cells (**Fig. 3G, 3H**), suggesting that B7-H3 promotes cell adhesion to ECM via control of PTGFRN/CD315, E-cadherin, integrin α3 and integrin αV.

**Fig. 3.**
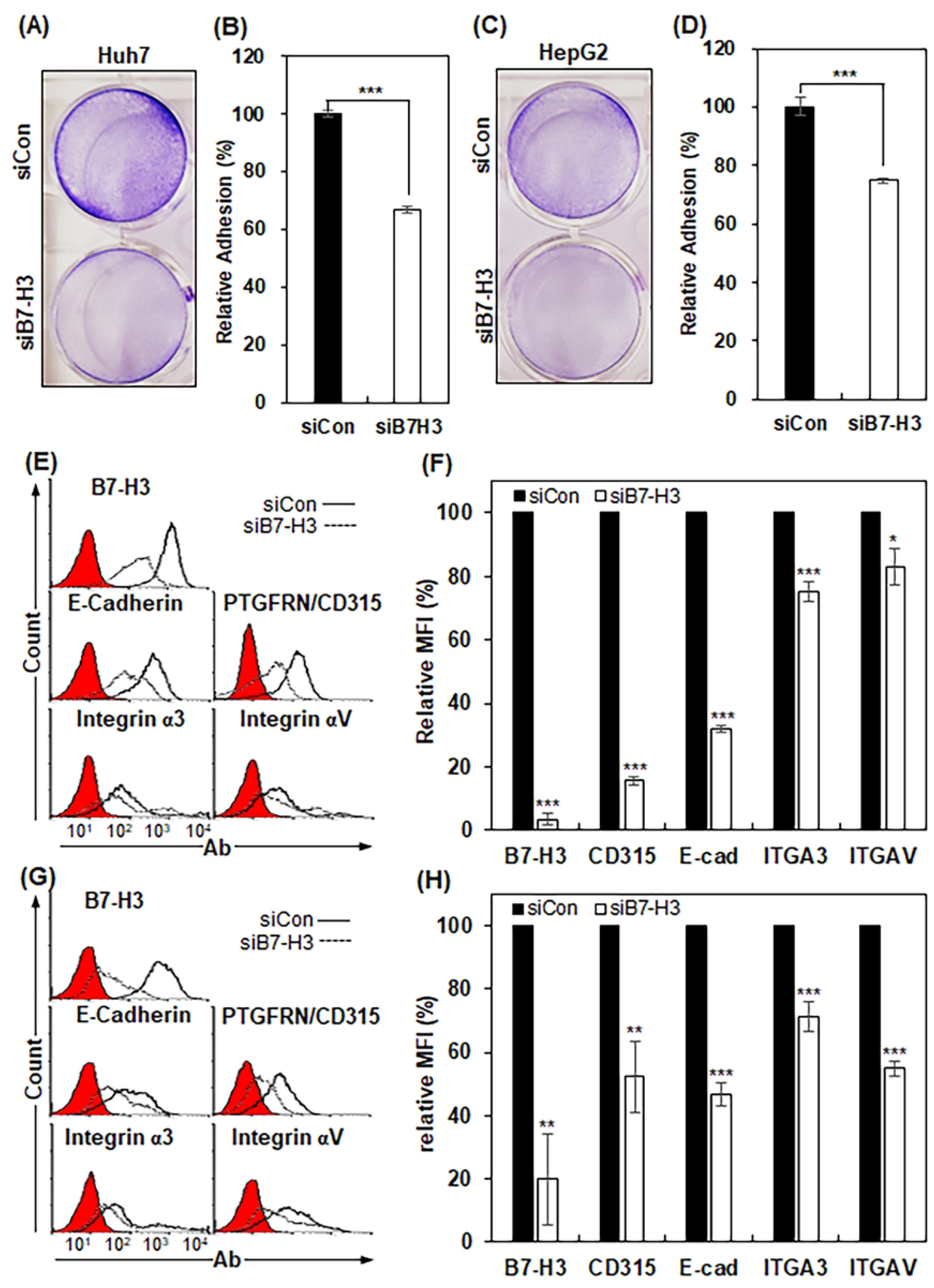
B7-H3 promotes cell adhesion in Huh7 and HepG2 cells. (A, C) Cell adhesion assays of B7-H3–knockdown Huh7 (A) and HepG2 (C) cells on gelatin-coated plates. Adherent cells were visualized by crystal violet staining. (B, D) Quantification of cell adhesion shown in (A) and (C), respectively. Data are presented as relative adhesion (%) compared with control siRNA (siCon)–transfected cells. (E, G) Flow cytometric analysis of cell adhesion molecule expression in B7-H3–knockdown Huh7 (E) and HepG2 (G) cells. (F, H) Quantification of mean fluorescence intensity (MFI) shown in (E) and (G), respectively, expressed as relative MFI (%) compared with siCon cells. *, p < 0.05; **, p < 0.01; ***, p < 0.005.

To further validate the functional role of B7-H3 in mesenchymal HCC cells, cell adhesion assays were also performed with *B7-H3* knockdown SNU449 cells. Adherent SNU449 cells were decreased by 27% in *B7-H3*-depleted SNU449 cells (**Fig. 4A, 4B**). Furthermore, in cell spreading assays, spreading of *B7-H3* KO cells was decreased by approximately 1.7-fold in Matrigel-coated plates in SNU449 cells (**Fig. 4C, 4D**). Similar results were also obtained with gelatin- and collagen IV-coated plates (**S2A-S2D Fig**), suggesting that B7-H3 promotes cell adhesion and spreading of HCC cells even in low density cultures. To examine whether *B7-H3* play a role in cell-to-cell adhesion, *B7-H3* KO cells were subjected to cell-to-cell adhesion assays on ultra-low attachment plates. Cell-to-cell aggregation was decreased by approximately 2-fold in *B7-H3* KO SNU449 cells (**Fig. 4E, 4F**). Quantitative PCR analysis showed that *B7-H3* KO decreased the expression of PTGFRN (CD315), E-cadherin, integrin αV (ITGAV) and integrin α3 (ITGA3) by approximately 73%, 77%, 67% and 95%, respectively, in SNU449 cells (**S2E Fig**). Western blotting also revealed that *B7-H3* KO decreased the expression of PTGFRN (CD315), E-cadherin, ITGAV and ITGA3 by 25%, 96%, 51% and 38%, respectively, in SNU449 cells (**Fig. 4G**). Taken together, the results suggest that B7-H3 promotes cell-to-ECM adhesion in HCC cells and also enhances cell-to-cell adhesion in mesenchymal HCC cells through control of the cell adhesion molecules.

**Fig. 4.**
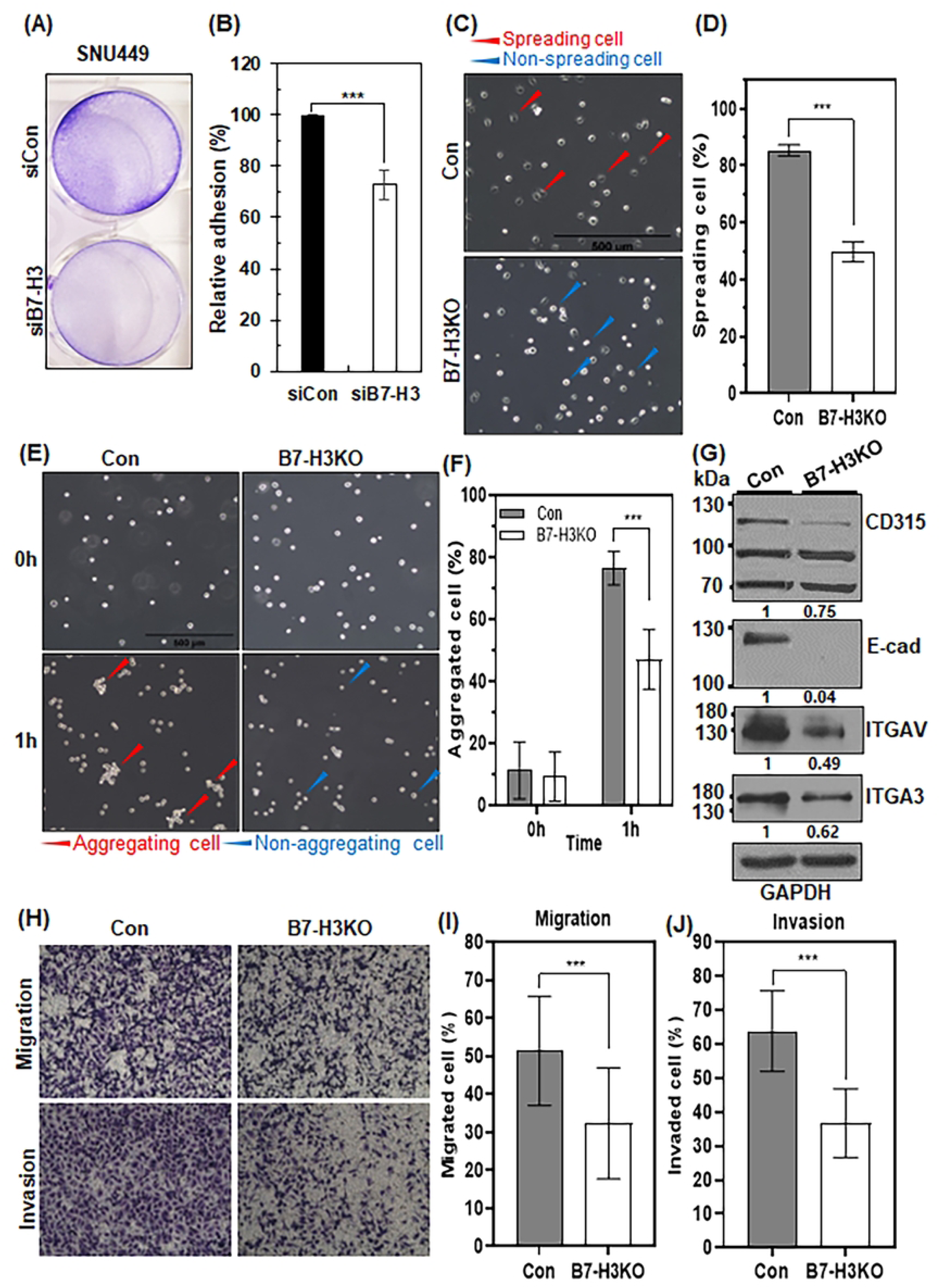
B7-H3 promotes cell adhesion, spreading, migration, and invasion in SNU449 cells. (A) Cell adhesion assay of B7-H3–knockdown SNU449 cells on gelatin-coated plates, visualized by crystal violet staining. (B) Quantification of adherent cells shown in (A), expressed as relative adhesion (%) compared with siCon cells. (C) Cell spreading assay of control and B7-H3 KO SNU449 cells on Matrigel-coated plates. (D) Quantification of spreading cells shown in (C). (E) Cell–cell aggregation assay of control and B7-H3 KO SNU449 cells cultured on ultra–low attachment plates. (F) Quantification of cell aggregation shown in (E). (G) Western blot analysis of E-cadherin, PTGFRN, ITGA3, and ITGAV in control and B7-H3 KO SNU449 cells. Protein levels were quantified using ImageJ software with GAPDH as a loading control. (H) Transwell migration and invasion assays of control and B7-H3 KO SNU449 cells. Migrated or invaded cells were stained with crystal violet. (I, J) Quantification of migrated (I) and invaded (J) cells shown in (H). ***, p < 0.005.

To take a look at whether B7-H3 is associated with the migratory potential of HCC cells, *B7-H3* KO SNU449 cells were subjected to transwell-migration assays. *B7-H3* KO decreased the migratory ability of SNU449 cells by approximately 19% (**Fig. 4H, 4I**). *B7-H3* KO also decreased the invasive ability of SNU449 cells by approximately 27% (**Fig. 4H, 4J**). The results suggest that B7-H3 positively regulates the migratory and invasive ability of HCC cells.

### B7-H3 supports expression of immune checkpoint molecules and suppresses NK cell-mediated cytotoxicity in HCC cells

B7-H3 is a member of the B7-family, including PD-L1 and PD-L2, and all of them act as immune checkpoint molecules in tumors [22]. Both B7-H3 and PD-L1 are regulated by miR-326 [23], and TGFβ1 also enhances both B7-H3 and PD-L1 in HCC cells [24], suggesting that they may be regulated simultaneously in HCC cells. To investigate whether B7-H3 regulates other immune checkpoint molecules, such as PD-L1, PD-L2, CD47 and CD24, on HCC cells, we examined the expression of immune checkpoint molecules on *B7-H3*-depleted HCC cells. Surface expression of B7-H3 was decreased in *B7-H3*-depleted Huh7 and HepG2 cells by approximately 96% and 80%, respectively (**Fig. 5A, 5B**). Under this condition, surface expression of PD-L1 and CD47 was decreased in *B7-H3*-depleted Huh7 cells by approximately 46% and 67%, respectively, whereas surface expression of CD24 was not altered. Surface expression of PD-L1 and CD47 was also decreased in *B7-H3*-depleted HepG2 cells by approximately 57% and 62%, respectively, whereas surface expression of CD24 was not affected (**Fig. 5C, 5D**). Surface expression of PD-L2 was not detected at all in Huh7 cells, while surface expression of PD-L2 was observed in HepG2 and was decreased by approximately 57% in *B7-H3*-depleted HepG2 cells.

**Fig. 5.**
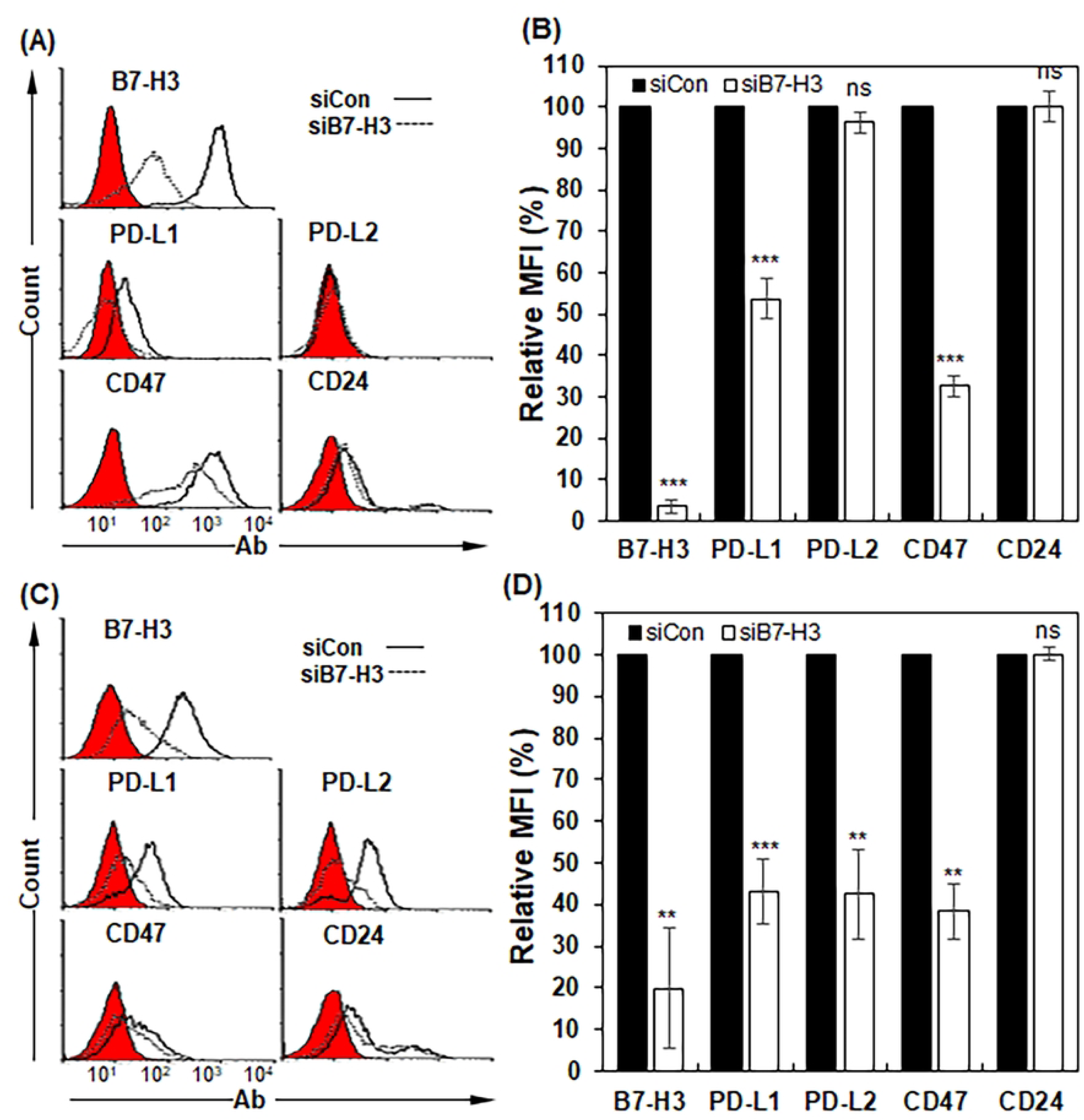
B7-H3 knockdown downregulates immune checkpoint molecules in Huh7 and HepG2 cells. (A, C) Flow cytometric analysis of PD-L1, PD-L2, CD47, and CD24 expression in B7-H3–knockdown Huh7 (A) and HepG2 (C) cells. (B, D) Quantification of immune checkpoint molecule expression shown in (A) and (C), respectively.

Surface expression of NPB40- and α-B7-H3-reactive B7-H3 was also decreased in *B7-H3*-depleted SNU449 cells by approximately 93% and 92%, respectively (**Fig. 6A, 6B**). Under this condition, surface expression of PD-L1, PD-L2 and CD47 was also decreased in *B7-H3*-depleted SNU449 cells by approximately 46%, 38% and 21%, respectively. Thus, cell surface expression of B7-H3 positively regulates surface expression of PD-L1, PD-L2 and CD47 in HCC cells. To further examine the relationship between B7-H3 and the immune checkpoint molecules, surface expression of PD-L1, PD-L2 and CD47 was also analyzed in *B7-H3* KO SNU449 cells. Surface expression of NPB40-reactive B7-H3 was decreased in *B7-H3* KO SNU449 cells by approximately 96% (**Fig. 6C, 6D**). Surface expression of PD-L1, PD-L2, and CD47 was also decreased in *B7-H3* KO SNU-449 cells by approximately 79%, 46% and 47%, respectively. Western blotting also showed the decreased expression of PD-L1, PD-L2 and CD47 in *B7-H3* KO SNU449 cells (**Fig. 6E**). The results suggest that B7-H3 expression is positively corelated to the expression of immune checkpoint molecules, such as PD-L1, PD-L2 and CD47.

**Fig. 6.**
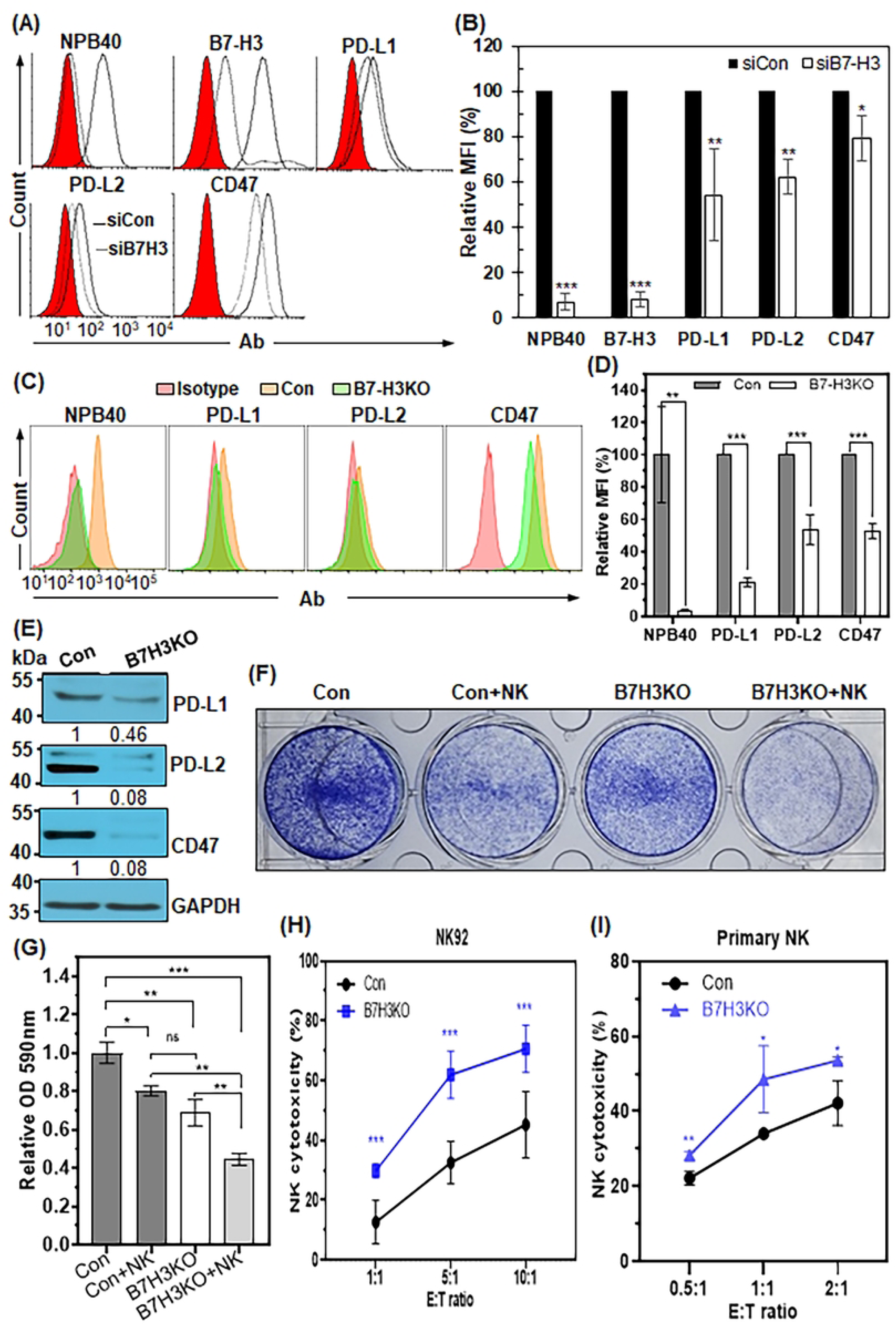
B7-H3 depletion downregulates immune checkpoint molecules and enhances NK cell–mediated cytotoxicity in SNU449 cells. (A) Flow cytometric analysis of PD-L1, PD-L2, CD47, and CD24 expression in B7-H3–knockdown SNU449 cells. (B) Quantification of MFI shown in (A), expressed as relative MFI (%) compared with siCon cells. (C) Flow cytometric analysis of PD-L1, PD-L2, and CD47 expression in control and B7-H3 KO SNU449 cells. (D) Quantification of MFI shown in (C), expressed as relative MFI (%) compared with control cells. (E) Western blot analysis of immune checkpoint proteins in control and B7-H3 KO SNU449 cells. Protein levels were quantified using ImageJ software with GAPDH as a loading control. (F) NK92 cell–mediated cytotoxicity assay. Control and B7-H3 KO SNU449 cells were co-cultured with NK92 cells at a 1:1 effector-to-target (E:T) ratio for 48 hours, followed by crystal violet staining of surviving tumor cells. Representative images from three independent experiments are shown. (G) Quantification of surviving cells shown in (F) by measuring absorbance at 590 nm. (H) NK92 cell–mediated cytotoxicity assessed by calcein-AM release assay at various E:T ratios. (I) Primary NK cell–mediated cytotoxicity assessed by calcein-AM release assay at various E:T ratios. ns, not significant; *, p < 0.05; **, p < 0.01; ***, p < 0.005.

To investigate whether *B7-H3* KO enhances NK cell-mediated cellular cytotoxicity in HCC cells, we applied two different assays to monitor NK cell mediated cytotoxicity: colony forming assay and calcein-AM assay. We cultured *B7-H3* KO SNU449 cells for 2 days with or without NK92 cells at a 1:1 NK:target cell ratio and measured the induction of cell death by staining the remaining SNU449 cells with crystal violet. Cellular cytotoxicity was increased when NK92 cells were added to SNU449 cells (**Fig. 6F, 6G**). When *B7-H3* KO SNU449 cells were treated with NK92 cells, NK92 cell-mediated cytotoxicity was further increased compared to when SNU449 cells were treated with NK92 cells (**Fig. 6F, 6G**). The calcein AM assay also showed that *B7-H3* KO SNU449 cells were more sensitive to NK92 cell-mediated cellular cytotoxicity than control cells at all E/T ratios (**Fig. 6H**). To further determine whether NK cell-mediated cytotoxicity is dependent on B7-H3 expression, *B7-H3* KO SNU449 cells were subjected to the same calcein AM cytotoxicity assays with primary NK cells. Primary NK cell-mediated cytotoxicity was also increased in *B7-H3* KO cells compared to control cells even at 0.5:1 E/T ratio (**Fig. 6I**). These findings indicate that B7-H3 expression leads to decreased NK cell-mediated cytotoxicity in HCC cells, and B7-H3 is a negative regulator of NK cell activity in HCC cells.

### B7-H3 function is mediated through Akt, MVP/ERK and FAK signaling in epithelial HCC cells and through FAK and JAK/STAT signaling in mesenchymal HCC cells

Previous studies showed that B7-H3 functions upstream from signal transduction pathways, such as the JAK2/STAT3, Akt/mTOR and MVP/ERK pathways, to induce proliferative, anti-apoptotic and metastatic mechanisms in cancer cells [10]. To figure out how B7-H3 promotes cancer progression in HCC cells, we analyzed signaling molecules in three HCC cell lines, Huh7, HepG2 and SNU449. Phosphorylation of Akt1/2/3 and ERK1/2 was decreased in *B7-H3* knockdown Huh7 cells by approximately 34% and 74%, respectively **(Fig. 7A, left panels**). Phosphorylation of S6, a downstream effector of Akt, was also decreased in *B7-H3* knockdown Huh7 cells by approximately 35% (**Fig. 7B, left panels**). MVP protein expression was also decreased by approximately 90% in *B7-H3* knockdown Huh7 cells (**Fig. 7B, left panel**). Cell surface MVP (csMVP) was also decreased in *B7-H3* knockdown Huh7 cells (**S3A, S3B Fig**). Similar results were also observed in HepG2 cells (**Fig. 7A, 7B, right panels**). Phosphorylation of FAK, a key regulator of adhesion and motility in cancer cells, was also decreased in *B7-H3* known Huh7 and HepG2 cells by approximately 78% and 96%, respectively (**Fig. 7A**). The results were expected because B7-H3 interacts with MVP in breast cancer cells and activates the ERK1/2 signaling pathway [25], and the function of MVP is also mediated through the FAK signaling in HCC cells [19]. Thus, the results suggest that the function of B7-H3 is mediated in epithelial HCC cells through the Akt1/2/3, MVP/ERK1/2 and FAK signaling pathways, which are critical for cancer growth, survival and metastasis.

**Fig. 7.**
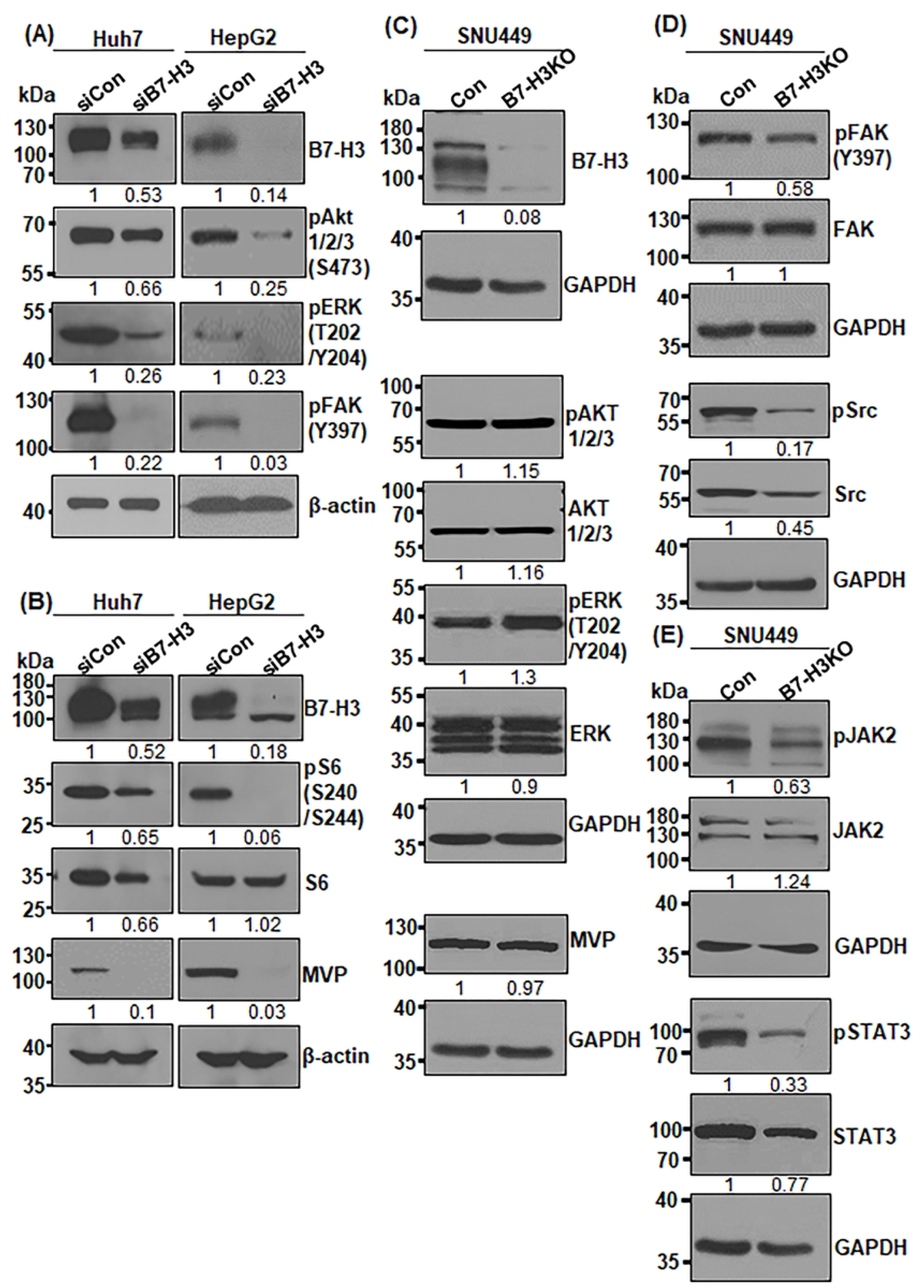
B7-H3 signaling is mediated through Akt/MVP/ERK and FAK/Src pathways in epithelial HCC cells and through JAK/STAT and FAK/Src pathways in mesenchymal HCC cells. (A, B) Western blot analysis of Akt1/2/3, ERK1/2, FAK, MVP, and S6 in B7-H3–knockdown Huh7 and HepG2 cells. Protein levels were quantified using ImageJ software with β-actin as a loading control. (C) Western blot analysis of Akt1/2/3, ERK1/2, and MVP in control and B7-H3 KO SNU449 cells. (D) Western blot analysis of FAK and Src in control and B7-H3 KO SNU449 cells. (E) Western blot analysis of JAK2 and STAT3 in control and B7-H3 KO SNU449 cells. For (C–E), GAPDH was used as a loading control.

To further examine B7-H3-associated signaling pathways in mesenchymal HCC cells, we analyzed signaling molecules in *B7-H3* KO SNU449 cells. Contrary to those epithelial HCC cells, interestingly, phosphorylation of Akt1/2/3 and ERK1/2 was slightly increased in *B7-H3* KO SNU449 cells (**Fig. 7C**). Furthermore, MVP protein expression was not significantly altered in *B7-H3* KO SNU449 cells (**Fig. 7C**), and csMVP expression was hardly detected in *B7-H3* knockdown and KO SNU449 cells (**S3C, S3D Fig**). The results suggest that MVP expression is not significantly associated with B7-H3 function in SNU449 cells. On the other hand, phosphorylation of FAK was decreased in *B7-H3* KO SNU449 cells by approximately 42% (**Fig. 7D**). Phosphorylation of Src, a downstream effector of FAK, was also decreased in *B7-H3* KO SNU449 cells by approximately 83% (**Fig. 7D**), which is consistent with those of *B7-H3* knockdown epithelial HCC cells. Phosphorylation of JAK2 and STAT3 was also decreased in *B7-H3* KO SNU449 cells by approximately 37% and 67%, respectively (**Fig. 7E**). Thus, the function of B7-H3 is mediated through the FAK/Src and JAK2/STAT3 signaling in mesenchymal HCC cells.

## Discussion

B7-H3 (CD276) is a promising pan-cancer antigen and a therapeutic target currently under investigation in numerous clinical trials [10, 26]. Our previous work demonstrated that an antibody (NPB40) targeting B7-H3 could inhibit tumor growth in an HCC xenograft model, highlighting its critical role in HCC progression [13]. The present study expands on these findings by comprehensively delineating the multifaceted mechanisms through which B7-H3 drives tumorigenesis in HCC cells. Our results show that B7-H3 is not only essential for cell survival, proliferation, migration and invasion but also coordinates cell adhesion and adhesion-related signaling. Furthermore, B7-H3 acts as a powerful immune evasion factor by suppressing NK cell-mediated cytotoxicity. These findings reveal previously unrecognized roles for B7-H3 in coordinating adhesion signaling and NK immune evasion in HCC

One of the key findings of this study is the critical role of B7-H3 in promoting cancer cell survival and inhibiting apoptosis. We observed that B7-H3 depletion, either transiently by siRNA or permanently by CRISPR/Cas9, significantly reduced clonogenic survival and enhanced apoptosis, particularly under anchorage-independent conditions (Fig. 1, 2). This is consistent with previous research showing B7-H3’s anti-apoptotic function in various cancers, often mediated by upregulating survival factors like Bcl-xL and downregulating pro-apoptotic proteins like Bax, as we found in Huh7 and HepG2 cells [27]. The ability of B7-H3 to promote cell survival under cellular stress is a crucial mechanism contributing to the aggressiveness of HCC. Our study further establishes B7-H3 as a major regulator of metastatic potential in HCC. We demonstrated that B7-H3 depletion significantly decreased cell adhesion, migration, and invasion (Fig. 4). This effect was correlated with the downregulation of several key cell adhesion molecules, including PTGFRN, E-cadherin, and integrins α3 and αV. This is particularly significant because B7-H3 expression has been shown to be prominent in tumor thrombi and intrahepatic metastases in HCC patients, suggesting its involvement in late-stage disease and metastasis [14]. Consistent with this notion, we found that B7-H3 captured approximately 56% (2501/4490) of all circulating tumor cells (CTCs) from patients with recurrent/metastatic HCC and metastatic liver cancers [28]. This was the second highest expression rate among the various markers we examined, following PTGFRN, suggesting that B7-H3 plays a significant role in CTCs from patients with advanced liver cancers [28]. The observation that B7-H3 KO also abolishes cell-to-cell aggregation and promotes apoptosis under detachment stress in mesenchymal SNU449 cells further supports its role in promoting cell aggregation and metastasis (Fig. 2C, 2D, 4E, 4F), potentially by facilitating the formation of circulating tumor cell (CTC) clusters, which are known to increase metastatic potential and resistance to apoptosis [29].

A particularly compelling finding is the role of B7-H3 in regulating the tumor immunosuppressive microenvironment. We discovered that B7-H3 expression is positively correlated with the surface expression of other key immune checkpoint molecules, including PD-L1, PD-L2, and CD47 (Fig. 5,6). This suggests that B7-H3 may function as a master regulator of immune evasion in HCC, coordinating multiple immunosuppressive pathways. The observation that B7-H3 depletion significantly enhances NK cell-mediated cytotoxicity is a novel and critical insight, underscoring its direct immunosuppressive role against NK cells, whose function in HCC is often compromised [30]. Given that B7-H3 is expressed in a much higher percentage of HCC patients (79-94%) compared to PD-L1 (25%), targeting B7-H3 may offer a broader and more effective immunotherapeutic strategy by simultaneously neutralizing multiple immunosuppressive pathways [7, 14].

Finally, our study provides a mechanistic understanding of B7-H3’s function by identifying distinct signaling pathways in different HCC subtypes. In epithelial HCC cells, B7-H3 promotes tumorigenesis through the Akt/S6, MVP/ERK, and FAK/Src pathways, consistent with reports of B7-H3-MVP interactions in other cancers [25]. Interestingly, in mesenchymal HCC cells, B7-H3’s effects were mediated primarily through the FAK/Src and JAK2/STAT3 pathways, with minimal involvement of the MVP/ERK axis. This suggests a context-dependent signaling role for B7-H3 and highlights the importance of considering HCC subtype when developing targeted therapies. This distinction explains why B7-H3 KO in mesenchymal cells could still lead to apoptosis despite a slight increase in ERK signaling, suggesting that the FAK/Src and JAK2/STAT3 pathways are more critical for survival in this phenotype. These insights are crucial for tailoring future anti-B7-H3 strategies and underscore the complexity of B7-H3 signaling in heterogeneous tumors.

In conclusion, our findings demonstrate that B7-H3 is a multifaceted driver of HCC progression, promoting survival, metastasis, and immune evasion through distinct, yet interconnected, signaling pathways. The ability of B7-H3 to regulate other immune checkpoints and directly suppress NK cell activity makes it a highly attractive target for novel immunotherapeutic approaches. Future research could focus on analyzing the impact of B7-H3 targeting on CTC clusters and exploring the specific interplay between B7-H3 and PTGFRN in immune regulation, further solidifying its potential as a superior therapeutic target in HCC.

## Supporting Information

**S1 Table. Antibodies used in this study.**

**S2 Table. Primers used for quantitative PCR analysis.**

**S1 Fig. Validation of B7-H3 knockdown and knockout in SNU449 cells.** (A) Western blot analysis of B7-H3 expression in control (siCon) and B7-H3–knockdown (siB7-H3) SNU449 cells. (B) Flow cytometric analysis of B7-H3 expression using NPB40 and anti–B7-H3 antibody (α–B7-H3). (C) Quantification of MFI shown in (B). (D) Western blot analysis of B7-H3 expression in control and B7-H3 KO SNU449 subclones. (E, F) Flow cytometric analysis of B7-H3 KO SNU449 subclones using NPB40 and quantification of relative MFI. (G, H) Western blot analysis of B7-H3 expression in control and representative B7-H3 KO subclone (B3). (I, J) Flow cytometric analysis and quantification of NPB40 binding in control and B7-H3 KO (B3) cells. ***, p < 0.005.

**S2 Fig. B7-H3 knockout reduces cell adhesion and downregulates adhesion molecule expression in SNU449 cells.** (A, B) Cell spreading assays of control and B7-H3 KO SNU449 cells on gelatin- (A) or collagen IV–coated plates (B). (C, D) Quantification of spreading cells shown in (A) and (B). (E) qPCR analysis of PTGFRN, E-cadherin, ITGAV, and ITGA3 expression in control and B7-H3 KO SNU449 cells. Data are presented as mean ± SD (n = 3). ***, p < 0.005.

**S3 Fig. MVP expression is associated with B7-H3 expression in Huh7 and HepG2 cells, but not in SNU449 cells**. **A,** Flow cytometric analysis of csMVP in B7-H3 knockdown Huh7 and HepG2 cells (siB7-H3). **B,** Statistics of A (**, p< .01; ***, p < .005). **C,** Flow cytometric analysis of csMVP in B7-H3 knockdown SNU449 cells. **D,** Flow cytometric analysis of csMVP in B7-H3 KO SNU449 cells.

## Acknowledgements

We thank Dr. Hee Chul Lee for his comments and proofreading of the manuscript. We express our deepest gratitude to our mentors Joo Yeo-Ja and Ryu Jeong-Rae, who have always provided the authors with undying motivation, inspiration, and passion. This study was supported in part by the National Research Foundation of Korea (RS2022-NR069561) and by the Korean Fund for Regenerative Medicine, funded by the Ministry of Science and ICT and the Ministry of Health and Welfare (21A0401L1). This research was also supported by the Regional Innovation System & Education (RISE) program through the Seoul RISE Center, funded by the Ministry of Education (MOE) and the Seoul Metropolitan Government (2025-RISE-01-019-04).

## Author Contributions

C.J. Ryu is the guarantor of this work and, as such, had full access to all the data in the study and takes responsibility for the integrity of the data and the accuracy of the data analysis. S.H. Han: development of methodology, acquisition of data, analysis and interpretation of data; Y.J Cheon: acquisition of data, development of methodology; H.M. Lee: acquisition of data, development of methodology; J.Y. Lee: acquisition of data; H. Seo: acquisition of data; M.J. Kim: resources, supervision; S.R. Yoon: resources, supervision; D. Choi: resources, supervision; All authors reviewed and approved the final manuscript.

## Ethics approval and consent to participate

Human UCB was obtained from healthy pregnant woman with consent and the study was conducted with the procedures approved by the Institutional Review Board (IRB) at Korea Research Institute of Bioscience and Biotechnology (IRB No. P-01–201610-31–002). All methods in this research were carried out in accordance with Declaration of Helsinki.

## Patient consent for publication

Not applicable

## Data availability

The datasets generated during and/or analyzed during the current study are available from the corresponding author on reasonable request.

